# Structural connectivity facilitates functional connectivity of auditory prediction error generation within a fronto-temporal network

**DOI:** 10.1101/365072

**Authors:** Lena K. L. Oestreich, Roshini Randeniya, Marta I. Garrido

## Abstract

Auditory prediction errors, i.e. the mismatch between predicted and actual auditory input, are generated by a hierarchical functional network of cortical sources. This network is also interconnected by auditory white matter pathways. Hence it would be reasonable to assume that these structural and functional networks are quantitatively related, which is what the present study set out to investigate. Specifically, whether structural connectivity of auditory white matter pathways enables effective connectivity of auditory prediction error generation. Eighty-nine participants underwent diffusion weighted magnetic resonance imaging. Anatomically-constrained tractography was used to extract auditory white matter pathways, namely the bilateral arcuate fasciculus, the inferior occipito-frontal fasciculi (IOFF), and the auditory interhemispheric pathway, from which Apparent Fibre Density (AFD) was calculated. The same participants also underwent a stochastic oddball paradigm, which was used to elicit prediction error responses, while undergoing electroencephalographic recordings. Dynamic causal modelling (DCM) was used to investigate the effective connectivity of auditory prediction error generation in brain regions interconnected by the above mentioned auditory white matter pathways. Brain areas interconnected by all auditory white matter pathways best explained the dynamics of auditory prediction error responses. Furthermore, AFD in the right IOFF and right arcuate fasciculus significantly predicted the effective connectivity parameters underlying auditory prediction error generation. In conclusion, the generation of auditory prediction errors within an effectively connected, fronto-temporal network was found to be facilitated by the structural connectivity of auditory white matter pathways. These findings build upon the notion that structural connectivity facilitates dynamic interactions within brain regions that are effectively connected.

**Significance statement:** The brain continuously generates and updates hypotheses that predict forthcoming sensory input. Within the auditory domain, it has repeatedly been reported that these predictions about the auditory environment are facilitated by specific functional cortical connections. These functionally connected brain regions are also structurally connected via auditory white matter pathways. For the first time, this study provides quantitative evidence for a structural basis along which this functional network of auditory prediction error generation operates. This finding provides evidence for the notion that the functional connectivity of dynamically interacting brain areas is facilitated by structural connectivity amongst these brain areas.

## Introduction

Auditory prediction errors are elicited by unexpected auditory events and result from a mismatch between predictions about forthcoming auditory sensations (which are based on past experiences), and actual auditory input (Friston, 2005). Previous studies used dynamic causal modelling (DCM) to investigate the effective connectivity of auditory prediction errors and identified a three-level hierarchical network comprised of connections from bilateral primary auditory cortex (A1) to the planum temporale (PT), and from the PT to the right inferior frontal gyrus (IFG) (Garrido et al., 2007; Garrido et al., 2008). This network included intrinsic (within source) connections within A1 and extrinsic (between-sources), forward and backward, connections between A1, PT and IFG. Interestingly, these effectively connected brain areas are also structurally connected via the auditory white matter pathways of the arcuate fasciculus, inferior occipito-frontal fasciculus (IOFF), and auditory interhemispheric pathway. It therefore seems feasible that auditory prediction errors are facilitated through dynamic interactions along these white matter tracts.

The arcuate fasciculus directly connects speech production areas in the IFG to auditory perception areas in the PT and also has shorter, indirect connections consisting of an anterior pathway which connects IFG to the inferior parietal lobule (IPL), and a posterior pathway which connects IPL to PT (Catani et al., 2005). While the posterior pathway has been reported to be involved in auditory comprehension, the anterior pathway is thought to be involved in the vocalization of semantic information (Catani et al., 2005). The IOFF connects areas in the frontal lobe (including IFG) with temporal areas (including PT and A1) and occipital areas (Martino et al., 2010) and has been reported to be involved in semantic language processing (Duffau et al., 2005). The auditory interhemispheric pathway is part of the corpus callosum, linking bilateral A1s and PTs and is thought to be involved in the integration of prosodic and syntactic information (Wigand et al., 2015).

Only few studies to date have attempted to directly associate measures derived from diffusion magnetic resonance imaging (MRI) with effective connectivity inferred from functional MRI (fMRI) data. A study combining DCM and diffusion weighted imaging (DWI) investigated cognitive control along the fronto-thalamic-cerebellar circuitry associated with symptoms in schizophrenia and found that fractional anisotropy in the anterior limb of the internal capsule was significantly correlated with fronto-thalamic effective connectivity in schizophrenia (Wagner et al., 2015). A study by Stephan et al. (2009) used probabilistic tractography from DWI to anatomically inform priors for DCM models. It was found that the DCMs that used anatomically informed priors outperformed DCMs without anatomically informed priors, which was taken to indicate that structural connectivity improves inferences about effective connectivity. Lastly, a study by Rae et al. (2015) correlated effective connectivity between IFG, pre-supplementary motor area (preSMA), subthalamic nucleus (STN) and primary motor cortex during an inhibitory control task with structural connectivity measures of white matter tracts connecting these regions and found a positive relationship of mean diffusivity and effective connectivity between preSMA and STN (Rae et al., 2015). While these studies provide evidence for an association between effective and structural connectivity, they fMRI data to infer effective connectivity. However, to date, no study has investigated whether structural connectivity predicts effective connectivity inferred from electrophysiological data, which, due to its higher temporal resolution, is more suited to investigate whether structural connectivity enables fast brain dynamics, such as auditory prediction error generation (which occur approximately 100 to 250ms after stimulus onset).

Here, we set out to investigate the relationship between structural connectivity (from DWI) and effective connectivity from Electroencephalographic (EEG) data, in the context of auditory prediction error generation. It was hypothesized that the effective connectivity network underlying auditory prediction error generation would include connections along all three auditory white matter pathways of the bilateral arcuate fasciculus, IOFF as well as the auditory interhemispheric pathway. It was furthermore hypothesized that the effective connectivity underlying auditory prediction error generation could be predicted from structural connectivity of the auditory white matter pathways.

## Methods

### Participants

Eighty-nine healthy participants (18 – 63 years, *M* = 25.02, *SD* = 10.34, 92.3% right-handed, and 57% female) were recruited through the online recruitment system - SONA and a weekly newsletter distributed to staff and alumni at the University of Queensland, Australia. All participants gave written informed consent and were monetarily reimbursed for their time. The study was approved by the University of Queensland Research Ethics Committee.

### Procedure

In the first part of the experiment, participants completed a set of questions about their demographic details. In the second part of the experiment, participants were seated in a quiet, dimly lit room, where they underwent an electroencephalography (EEG) recording. The experimental design consisted of an auditory frequency oddball paradigm (Garrido et al., 2013a) and a simultaneous N-back task (Sweet, 2011). Participants listened to a stream of sounds with log-frequencies sampled from two Gaussian distributions with equal means (500Hz) and different standard deviations (narrow: *σ_n_* = .5 octaves; broad: *σ_b_* = 1.5 octaves). All tones were played with duration of 50ms, including 10ms smooth rise and fall periods and inter-stimulus intervals of 500ms. 10% of the tones were defined as standard tones, which were played at 500Hz, i.e. at the mean of both distributions, and 10% of the tones were defined as deviant tones, which were played at 2000Hz, i.e. at the tails of the distributions and hence outliers. Standard and deviant tones were inserted into the sound stream pseudo-randomly with the Gaussian contributing 80% of the tones. Participants were told to disregard the tones and to focus on the visual N-back task, in which they were instructed to press a button every time they saw the same letter was played 2 trials beforehand. The timings of the sounds and the visual N-back task were independent of each other in order to avoid motor and attention artefacts. The overall experiment lasted approximately 30 minutes and was divided into 4 blocks. The narrow and broad distribution conditions were presented in separate blocks and the order of the blocks was counter-balanced across participants.

In the third part of the experiment, participants underwent diffusion-weighted and T1-weighted magnetic resonance imaging (MRI) scans.

### Data acquisition and pre-processing Electroencephalography (EEG)

Continuous EEG was recorded with a 64 Ag/AgCl BioSemi ActiView system at a sampling rate of 1024Hz. External electrodes were placed on the outer canthi of both eyes, below and above the left eye and on both mastoids. Pre-processing and data analysis were performed using SPM12 (Wellcome Trust Centre for Neuroimaging, London; http://www.fil.ion.ucl.ac.uk/spm/) with MATLAB (MathWorks). Data were referenced to the common average reference and high-pass filtered above 0.5Hz. Eye blinks were detected and removed with the vertical electro-oculogram (VEOG) and horizontal electro-oculogram (HEOG) channels. The data were epoched offline into 500ms intervals with 100ms pre- and 400ms post-stimulus onset. Trials containing artefacts exceeding ±50µV were rejected. The remaining artefact free trials were robustly averaged to event-related potentials (ERPs), low-pass filtered below 40Hz and baseline corrected using the −100 to 0ms pre-stimulus interval.

### Diffusion-Weighted Imaging (DWI)

Two diffusion-weighted (DW) image series were acquired on a Siemens Trio 3T system (Erlangen, Germany) using an echo-planar imaging (EPI) sequence. The first DW image series consisted of a high *b*-value data set with field-of-view (FoV) of 220mm, phase partial Fourier (PPF) 6/8, parallel acceleration factor 2, 55 slices, 2mm isotropic resolution, 64 diffusion-sensitization directions at *b*=3000s/mm^2^ with two *b*=0 volumes, TR=8600ms, TE=116ms, and 10min acquisition time. The second DW image series consisted of a low *b-*value data set with FoV 220mm, PPF 6/8, parallel acceleration factor 2, 55 slices, 2mm isotropic resolution, 33 diffusion-sensitization directions at *b*=1000s/mm^2^ with one *b*=0 volumes, TR=8600ms, TE=116ms, and 5min acquisition time. A T1-weighted image data set was acquired with the magnetisation-prepared two rapid acquisition gradient echo (MP2RAGE) sequence (Marques et al., 2010) with FoV 240mm, 176 slices, 0.9mm isotropic resolution, TR=4000ms, TE=2.92ms, TI1=700ms, TI2=2220ms, first flip angle=6°, second flip angle=7°, and 5min acquisition time. Three *b*=0 images were acquired interspersed between the DW image series and the MPRAGE sequence, with reversal of the acquisition direction along the phase-encoded axis for one of the three images and acquisition time of 30ms each.

DW images were corrected for signal intensity inhomogeneities (Zhang et al., 2001) as well as head movements and eddy current distortions using the FSL TOPUP (Smith et al., 2004) and EDDY (Andersson and Sotiropoulos, 2016) tools. The remaining processing steps were conducted using tools implemented in MRtrix3 (Tournier et al., 2012). DW and T1-weighted images were co-registered using boundary-based registration (Greve and Fischl, 2009). A five-tissue-type segmented image (cortical grey matter, sub-cortical grey matter, white matter, cerebrospinal fluid, pathological tissue) was generated from the structural images pre-processed using the recon-all command in FeeSurfer (Dale et al., 1999). Response functions were estimated using the multi-shell, multi-tissue algorithm implemented in MRtrix3 (Jeurissen et al., 2014). Multi-tissue constrained spherical deconvolution was applied to obtain fiber orientation distributions (FOD; (Jeurissen et al., 2014). Anatomically-constrained tractography (ACT; (Smith et al., 2012)) was used to generate probabilistic streamlines of the auditory interhemispheric pathway, bilateral IOFF, and bilateral arcuate fasciculus with a maximum path length of 200mm, a minimum path length of 4mm, step size of 1mm and back-tracking (see Figure 3A). Spherical deconvolution Informed Filtering of Tractograms (SIFT2) is a filtering algorithm to remove inadequacies resulting from the reconstruction method to create tracts that are more biologically plausible (Smith et al., 2013). SIFT2 provides a cross-sectional area multiplier for each streamline, such that the contribution of each streamline to the tractogram can be weighted, whilst simultaneously retaining all reconstructed streamlines (Smith et al., 2015). Contrary to the traditional diffusion tensor model, which estimates average values across an entire voxel, the apparent fibre density (AFD) is a metric derived from the FOD lobe parallel to the direction of the streamline and therefore provides a sub-voxel, tract specific measure (Raffelt et al., 2012). The AFD integral for a specific FOD lobe in a specific direction is approximately proportional to the intra-axonal volume of the corresponding white matter fibre bundle oriented in that direction. AFD can therefore be defined as the fraction of space occupied by a white matter fibre bundle (Wright et al., 2017). The apparent fibre density (AFD) related to the total intra-axonal tract volume was calculated by summing the integrals for all FODs associated with the tract streamlines and dividing it by the streamline length (Raffelt et al., 2012).

### Source reconstruction and analysis

Source images were created from the scalp activity using a Boundary Element Method (BEM) and a standard MNI template for the cortical mesh. Reconstructed images were obtained for the conditions *Standard Narrow, Deviant Narrow, Standard Broad and Deviant Broad* in each participant and smoothed at FWHM 8×8×8 mm^3^. A mixed analysis of variance (ANOVA) was conducted with the within-subjects factors *surprise* (standards/deviants) and *variance* (narrow/ broad) and significant main effects and interactions were followed-up with t-statistic contrasts. Effects are displayed at an uncorrected threshold of *p* < .05.

### Dynamic causal modelling (DCM)

DCM is based on a generative spatiotemporal model for EEG responses elicited by experimental stimuli (Kiebel et al., 2008). DCM uses neural mass models to make inferences about source activation as well as the dynamic interactions, or connectivity, amongst these sources (Jansen and Rit, 1995; David and Friston, 2003). In DCM, nodes are hierarchically organized and interconnected via forward, backward and lateral connections (Felleman and Van Essen, 1991; David et al., 2005; Kiebel et al., 2007). DCMs are defined to test theoretically informed hypotheses about several connectivity models which define alternative networks that explain (or predict) the generation of ERP signals (Garrido et al., 2008).

### Bayesian model selection and averaging

Bayesian Model Selection (BMS) was used to compare several plausible network connections by estimating the probability of the data given a particular model within the space of models compared (Penny et al., 2004). The winning model is the one, which displays the best balance between accuracy maximisation and complexity minimization.

The posterior probability of each model was calculated over all participants using a random effects approach (RFX; (Stephan et al., 2009), which quantifies the probability that a particular model generated the data for any randomly chosen participant. The reported exceedance probability is the probability that one model is more likely than any other model, given the group data (Stephan et al., 2010). The main conclusions of the family comparisons are based on inferences with RFX exceedance probabilities of .89 on average (ranging from .78-1). The Bayesian omnibus risk (BOR) directly measures the probability that all model frequencies are indistinguishable, which quantifies the risk incurred when performing Bayesian model selection (Rigoux et al., 2014). The BOR is defined by a value between 0 and 1, whereby a value close to 1 indicates that the models are indistinguishable and a value close to 0 indicates that the models are well distinguishable from each other. In order to make inferences on individual connections, we used random-effects Bayesian Model Averaging (BMA) which computes a weighted average estimate of the individual connections across all models and participants, such that each model contributes to the overall estimate according to its posterior probability (Stephan et al., 2009).

### DCM specification

The models compared in the present study include up to 10 bilateral brain regions, which were hierarchically organized into four levels. These alternative models were motivated by brain regions, which have previously been shown to be implicated in auditory prediction errors (Garrido et al., 2013b), appeared in the source reconstruction of the present data and are interconnected by the auditory white matter pathways of the arcuate fasciculus, the auditory interhemispheric pathway and the inferior occipito-frontal fasciculus (IOFF). The bilateral primary auditory cortex (A1) was defined as the input node of auditory information. The arcuate fasciculus comprises a direct pathway between planum temporale (PT) and inferior frontal gyrus (IFG) as well as two indirect pathways, namely the anterior pathway, which connects inferior parietal lobule (IPL) with IFG and the posterior pathway, which connects PT with IPL. The IOFF interconnects IFG with PT and the occipital lobe (OL). Lastly, the auditory interhemispheric pathway connects A1 to PT in the left hemisphere to A1 and PT in the right hemisphere. The coordinates were chosen based on previous studies on MMN generation for STG, IFG (Opitz et al., 2002) and A1 (Rademacher et al., 2001) and language processing for IPL (Bitan et al., 2005) and PT (Osnes et al., 2011). The mean locations for the nodes were based on the Montreal Neurological Institute (MNI) coordinates for left A1 (−52, −19, 7), right A1 (50, −21, 7), left PT (−57, −20, 1), right PT (54, −19, 1), left IPL (−53, −32, 33), right IPL (51, −33, 34), left IFG (−48, 13, 17), right IFG (49, 12, 17), left OL (−45, −75, 11) and right OL (44, −75, 5; see Figure 1 (Lacadie et al., 2008)).

A model space of 50 models was considered, including symmetric and non-symmetric hierarchical models, with and without interhemispheric connections between the left and right A1 and PT, with and without indirect connections between PT and IFG via IPL as well as models with and without connections to OL (for a full description of the model space see Figure 1 – note that 25 model architectures are displayed, which were tested with and without interhemispheric connections). All models included modulations of intrinsic connectivity at the level of A1 and were individually estimated and compared to each other using BMS. To investigate whether auditory prediction errors operate along interhemispheric connections in addition to unilateral connections, a family including all models with interhemispheric connections (i.e. *lateral connections* family) was compared to a family without interhemispheric connections (i.e. *no lateral connections* family). Lastly, to investigate whether connections along individual white matter tracts or a combination of all tracts were more likely to be utilized during auditory prediction error generation, the models were partitioned into four different families, namely the *auditory interhemispheric pathway* family, the *IOFF* family, the *arcuate fasciculus* family, and the *auditory interhemispheric pathway + IOFF + arcuate fasciculus* family (see Figure 1). Each of the 50 models was fitted to every participant’s mean ERP response in the time window of 0-400ms after stimulus onset. The *Standard Narrow* condition elicited the lowest ERP amplitude in the MMN time window (100-200ms after stimulus onset) and was used as the baseline condition (weight = 0), followed by increasing ERP amplitudes in the *Standard Broad* (weight = 1) *Deviant Broad* (weight = 2) and *Deviant Narrow* (weight = 3) conditions (see Figure 2A).

**Figure 1.**
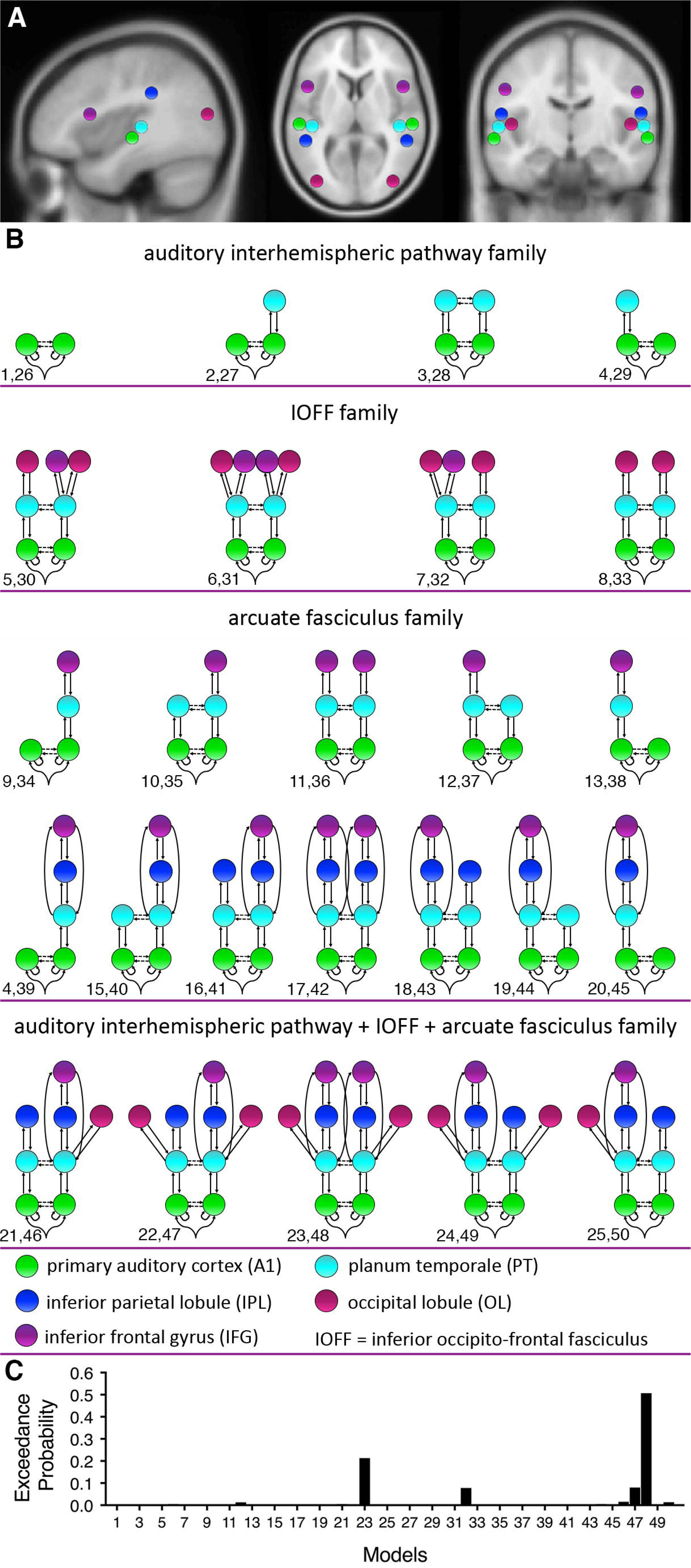
A) Prior mean locations for the DCM nodes and model space. The MNI coordinates include for the prior mean node locations: left A1 (−52, −19, 7), right A1 (50, −21, 7), left PT (−57, −20, 1), right PT (54, −19, 1), left IPL (−53, −32, 33), right IPL (51, −33, 34), left IFG (−48, 13, 17), right IFG (49, 12, 17), left OL (−45, −75, 11) and right OL (44, −75, 5). B) Model space. Models 1-25 excluded interhemispheric connections between the A1s and PTs and models 26-50 included interhemispheric connections between the A1s and PTs. These 50 models were defined to test different hypotheses about the effective anatomy of auditory prediction error generation. The models were combined to 4 families including the *auditory interhemispheric pathway* family (model 1-4 and 26-29), the *IOFF* family (models 5-8 and 30-33), the *arcuate fasciculus* family (models 9-20 and 34-45), and *the auditory interhemispheric pathway + IOFF + arcuate fasciculus* family (models 21-25 and 46-50). C) Model exceedance probability for auditory prediction error generation. Bayesian model selection (random effects) over the whole model space indicated that auditory prediction error generation was best explained by a model with intrinsic connections within A1, interhemispheric connections between left and right A1 and interhemispheric connections between left and right PT, as well as recurrent (forward and backward) connections between A1, PT, IPL, OL, IPL and IFG (model number 48).

**Figure 2.**
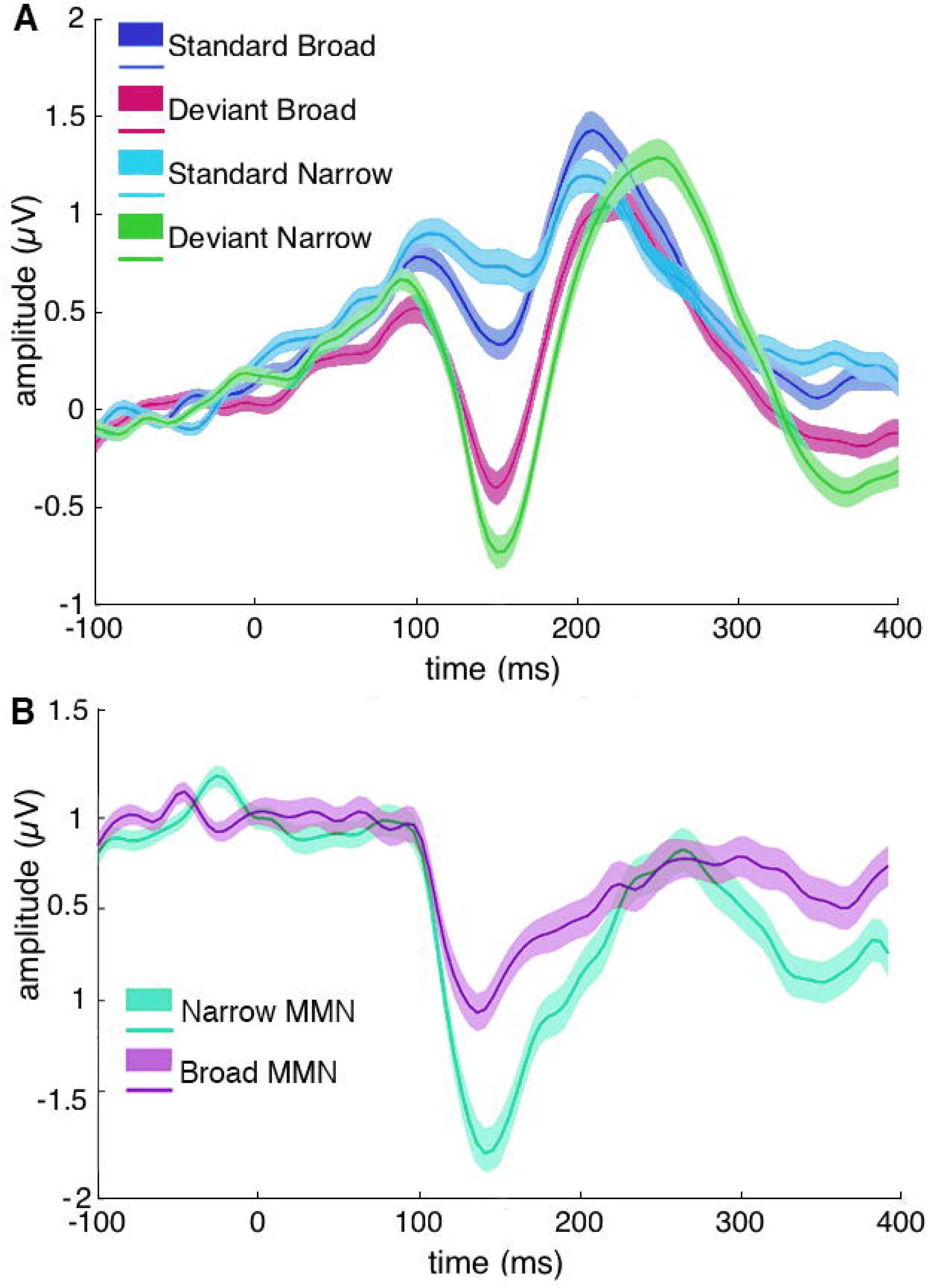
Event-related potentials (ERPs) elicited by A) the *Standard Narrow* (light blue), *Standard Broad* (dark blue)*, Deviant Broad* (magenta), and *Deviant Narrow* (green) conditions and B) mismatch negativity (MMN) waveforms (difference between responses to deviant and standard tones) for the broad (purple) and narrow (cyan) conditions extracted from electrode Fz. ERPs are time-locked to the onset of tones. Solid lines indicate the mean and lighter shading indicates standard error of the mean.

### Predicting effective connectivity from structural connectivity

A multivariate normal regression implemented in Matlab (Meng and Rubin, 1993; Little and Rubin, 2002) was fit to the DCM connectivity parameters derived from the BMA (26 outcome variables: intrinsic connections within left and right A1, interhemispheric connections between left and right A1, and left and right STG; reciprocal – forward and backward - connections linking A1 and PT, PT and IPL, IPL and IFG, PT and IFG, as well as PT and OL) where the effect of AFD was tested (5 independent variables: AFD of the right arcuate fasciculus, left arcuate fasciculus, right IOFF, left IOFF and the auditory interhemispheric pathway). In case of significant contributions of individual white matter pathways in predicting effective connectivity of auditory prediction error generation across the whole network, the association between the structural connectivity of those individual white matter pathways and their respective effective connectivity parameters was tested. For this purpose, multivariate linear regressions were performed with the AFD of the individual pathways as outcome variables and the corresponding DCM connectivity parameters defined along those pathways as predictor variables. Bonferroni corrections were used to correct for multiple comparisons.

## Results

The event-related potential (ERP) waveforms of the prediction errors responses, derived by subtracting the standards from the deviants, are displayed for the broad and narrow conditions in Figure 2B.

### Source Level

Putative EEG sources were reconstructed from the scalp activity in order to infer the cortical sources most likely to have generated the observed EEG signal. A *surprise*variance* interaction effect was observed across right temporal (peak-level *F* = 9.44, *p_uncorr_* = .002), left occipital (peak-level *F* = 8.43, *p_uncorr_* = .004) and central (peak-level *F* = 5.32, *p_uncorr_* = .021) regions and a main effect of *surprise* was observed across left frontal (peak-level *F* = 5.09, *p_uncorr_* = .024), right frontal (peak-level *F* = 4.66, *p_uncorr_* = .031) and left occipital regions (peak-level *F* = 5.11, *p_uncorr_* = .024). These source activations are in line with previous findings in studies using MMN paradigms (Doeller et al., 2003; Garrido et al., 2013b) and are interconnected by auditory white matter pathways (see Figure 3A).

**Figure 3.**
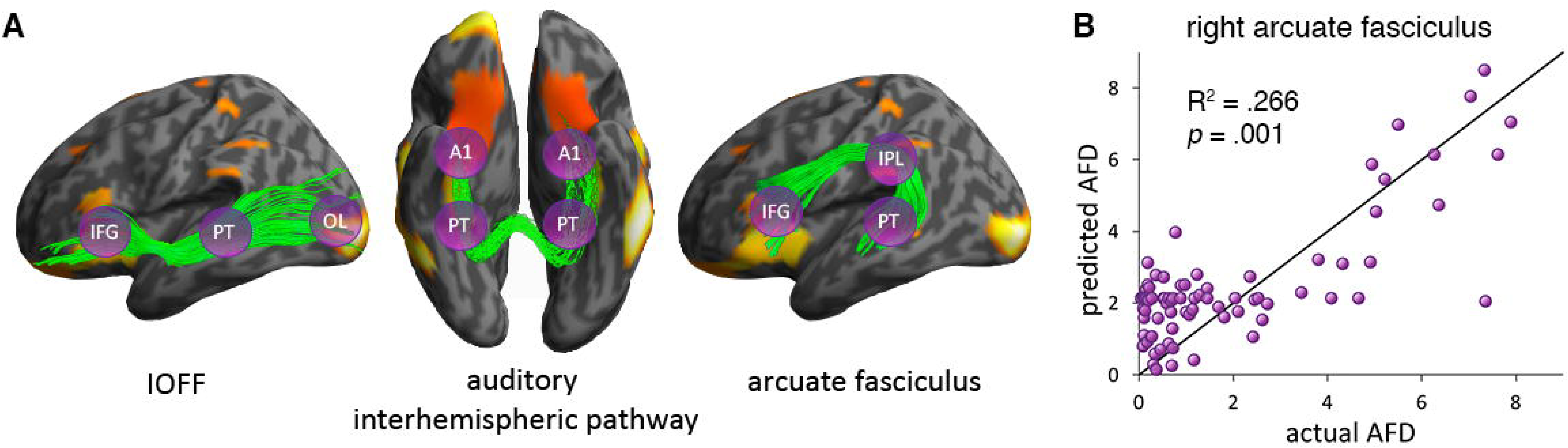
A) Auditory white matter pathways overlaid on source reconstructed images for the interaction between surprise and variance. The inferior occipito-frontal fasciculus (IOFF) connects IFG, PT and OL (left), the auditory interhemispheric pathway connects bilateral A1s and PTs (middle) and the arcuate fasciculus connects IFG, IPL and PT (right). B) Regression analysis. Observed apparent fibre density (AFD) plotted against the predicted AFD of the right arcuate fasciculus.

### DCM analysis

In a first step, all 50 models with and without interhemispheric connections between bilateral A1 and PT were individually compared to each other. Results indicated that the best model included bilateral forward and backward connections between A1 and PT, PT and IPL, PT and OL, IPL and IFG, direct connections between PT and IFG as well as interhemispheric connections between left and right A1 and left and right PT (exceedance probability = .51; BOR = 9×10^−6^; see Figure 2C).

In order to investigate whether auditory prediction errors engage interhemispheric connections in addition to unilateral connections, a family including all models with interhemispheric connections was compared to a family without interhemispheric connections. BMS revealed that the *lateral connections* family was more likely than the *no lateral connections* family (exceedance probability = .78).

To test for specific hypotheses as to whether connections along individual white matter pathways or a combination of connections along all pathways are more likely to be engaged during prediction error generation, the models were partitioned into four different families as described in the method section (see Figure 1B). BMS of these families indicated that the *auditory interhemispheric + IOFF + arcuate fasciculus* family was the winning family (exceedance probability = .86), indicating that effective connectivity of auditory prediction errors is best explained by regions that are interconnected by all three auditory white matter pathways as opposed to one particular tract.

### Predicting effective connectivity from structural connectivity

Six statistical outliers (3 in the right arcuate fasciculus, one in the auditory interhemispheric pathway and 2 in the left arcuate fasciculus), defined as having AFD values greater than 3 standard deviations from the mean were excluded from further analyses.

A multivariate normal regression was conducted in order to investigate whether the effective connectivity that underpins auditory prediction error generation could be predicted based on the structural connectivity of auditory white matter pathways. This was done using the AFD values from the bilateral IOFF, arcuate fasciculus, as well as the auditory interhemispheric pathway, as predictors and the twenty-six connectivity parameters derived with BMA over all DCMs as outcome variables. The structural connectivity of the right IOFF (95% CI [.000048, .000254]) and the right arcuate fasciculus (95% CI [.000346, .006974]) showed significant associations with the effective connectivity of the whole network, indicating that effective connectivity of auditory prediction error generation increases with increasing structural connectivity (i.e. AFD) in the right IOFF and arcuate fasciculus.

In order to follow up these association, we ran two multivariate linear regression analyses. In the first regression analysis, AFD of the right IOFF was entered as the outcome variable and the DCM connectivity parameters defined along the IOFF as the independent variables (i.e., PT to IFG, IFG to PT, PT to OL, and OL to PT). The overall model explained 12% of the variance of AFD of the right IOFF but did not remain significant after Bonferroni correction (R^2^ = .120, Bonferroni corrected *p* = .078). In the second regression analysis, AFD of the right arcuate fasciculus was entered as the outcome variable and the DCM connectivity parameters defined along the arcuate fasciculus as the independent variables (i.e. PT to IFG, IFG to PT, IPL to IFG, IFG to IPL, PT to IPL, IPL to PT). The overall model explained 27% of the variance of AFD of the right arcuate fasciculus (R^2^ = .266, Bonferroni corrected *p* = .001; see Figure 3B). Specifically, the AFD of the right arcuate fasciculus was trend-level predicted by the forward connection from PT to IFG (95% CI [−.149, 11.880]) and significantly predicted by the forward connection from PT to IPL (95% CI [.858, 9.028]), the backward connection from IPL to PT (95% CI [−20.053, −2.534]), and the backward connection from IFG to IPL (95% CI [5.789, 19.360]). This indicates that the structural connectivity along the arcuate fasciculus predicts the effective connectivity amongst the cortical regions that lie within it, namely PT, IPL and IFG.

## Discussion

The first aim of this study was to explore whether brain regions that are effectively connected during auditory prediction error generation operate along auditory white matter pathways. As hypothesized, the winning DCM model comprised brain regions interconnected by all included auditory white matter pathways, namely the bilateral arcuate fasciculus, bilateral IOFF and auditory interhemispheric pathway. The second aim of this study was to investigate whether the underlying effective connectivity could be predicted from the structural connectivity. We found that the white matter microstructure of the right IOFF and right arcuate fasciculus were significant predictors of effective connectivity across the whole network. Moreover, the backward connections from the inferior parietal lobule to the planum temporale, and the inferior frontal gyrus to the inferior parietal lobule, as well as the forward connections from planum temporale to inferior parietal lobule individually predicted the microstructure of the right arcuate fasciculus. These findings indicate that auditory prediction errors are generated along a fronto-temporal auditory network that is both structurally and effectively connected.

In line with previous findings, the results of the present study indicate that auditory prediction errors are generated by hierarchically organised cortical brain regions including intrinsic connections and extrinsic forward and backward connections (Garrido et al., 2007; Garrido et al., 2008). Specifically, the model that best explained auditory prediction error generation included intrinsic connections at A1, interhemispheric connections between bilateral A1s and PTs, as well as forward and backward connections between A1 and PT, PT and IPL, PT and IFG, PT and OL, and IPL and IFG. The present study refined previously employed models, which included recurrent connections between A1, PT and IFG, by now including recurrent connections between PT and OL, PT and IPL, and IPL and IFG. These areas were included due to the fact that they are structurally interconnected by the auditory white matter pathways of the arcuate fasciculus, IOFF, and the auditory interhemispheric pathway. Furthermore, a study by (Garrido et al., 2013b) employed source reconstruction on MEG data using the same MMN paradigm as used in the present study and found that the OL and IPL were engaged in prediction error generation. It is therefore not surprising that the winning model of the present study outperforms those with a simpler architecture used in previous studies (Garrido et al., 2007; Garrido et al., 2008).

The current study is one of the few studies to integrate structural neuroimaging measures with electrophysiological brain function measures (Salisbury et al., 2007; Fusar-Poli et al., 2011; Whitford et al., 2011) and, to the best of our knowledge, the first to integrate structural connectivity with EEG derived connectivity, within the same sample. We observed that the microstructure of the right IOFF and the right arcuate fasciculus significantly predicted the effective connectivity underlying auditory prediction errors. Additionally, we found that the effective connectivity parameters along the right arcuate fasciculus accounted for a significant amount of variance (27%) of the right arcuate fasciculus microstructure. In line with previous studies (Garrido et al., 2007; Garrido et al., 2008), the current study also found that auditory prediction error generation is facilitated by an effective network of intrinsic connections within primary auditory cortex, forward and backward connections from bilateral primary auditory cortex to bilateral superior temporal gyrus (i.e. planum temporale, more specifically). Contrary to previous studies however, which reported reciprocal connections from the superior temporal gyrus (i.e. planum temporale) to the inferior frontal gyrus (Garrido et al., 2007; Garrido et al., 2008) on the right hemisphere only, we observed reciprocal connections from superior temporal gyrus to planum temporale bilaterally, which could be explained by the fundamentally different nature of the statistical regularity violation posed by the paradigm used here (Garrido et. al, 2013). Nevertheless, our finding of structural connectivity within the right IOFF and right arcuate fasciculus predicting effective connectivity of auditory prediction error generation supports the previous findings of a right lateralized network.

Most of the effective connectivity parameters showed a positive association with the AFD values of the corresponding white matter pathways, such that effective connectivity increased as structural connectivity. This indicates that strong microstructural integrity of the right arcuate fasciculus facilitates improved effective connectivity of auditory prediction error generation. However, the backward connection from inferior parietal lobule to planum temporale showed a negative association with the microstructure in the right arcuate fasciculus. This finding of decreased structural connectivity with increasing effective connectivity may seem counter intuitive. AFD is a measure of intra-axonal volume fraction (Raffelt et al., 2012), which can be interpreted such that high AFD values correspond to a high axonal density and low values indicate a reduction of axonal density. According to this interpretation, AFD values should increases with increasing effective connectivity, as opposed to decrease. One potential explanation for this finding is that since AFD is normalized by the total fibre length, participants with well-preserved longer-fibre communication and diverse branching might show decreased fibre density.

The winning DCM model included brain areas along all auditory white matter pathways. However, the regression model did not confirm individual significant contributions of structural connectivity of the auditory interhemispheric pathway for predicting effective connectivity engaged in the generation of auditory prediction errors. The interhemispheric pathway has mainly been associated with the integration of prosody and syntax (Wigand et al., 2015). The IOFF and the arcuate fasciculus on the other hand, have been implicated in auditory and language processing more generally (Catani and Thiebaut de Schotten, 2008). The paradigm used in the present study consisted of simple tones instead of speech sounds and participants were passively listening to tones, rather than generating sounds themselves. It is therefore possible that even though the winning DCM model included areas along all of the auditory white matter tracts, areas interconnected by the IOFF and particularly the arcuate fasciculus, play stronger roles in auditory prediction error generation than the auditory interhemispheric pathway.

In conclusion, we found that a functional network along the auditory white matter pathways of the bilateral arcuate fasciculus, bilateral IOFF and the auditory interhemispheric pathway best explained the generation of auditory prediction errors in a statistical oddball paradigm. Critically, we observed, for the first time, that the structural connectivity (i.e. AFD) within the right IOFF and the right arcuate fasciculus significantly predicted the effective connectivity of this functional network. These auditory white matter pathways interconnect dynamically interacting brain regions and therefore provide a structural basis along which auditory prediction error generation can functionally operate. Taken together, these findings indicate that in a stochastic environment, auditory prediction errors recruit brain regions that are effectively and structurally connected by auditory white matter pathways.

## Acknowledgements

This work was funded by the Australian Research Council Centre of Excellence for Integrative Brain Function (ARC Centre Grant CE140100007), a University of Queensland Fellowship (2016000071), and a Foundation Research Excellence Award (2016001844) to MIG, as well as a University of Queensland International Research Scholarship to RR. The authors thank the participants for their time and Aiman Al-Najjar and Nicole Atcheson for assisting with data collection.

